# Efficient estimation and applications of cross-validated genetic predictions

**DOI:** 10.1101/517821

**Authors:** Joel Mefford, Danny Park, Zhili Zheng, Arthur Ko, Mika Ala-Korpela, Markku Laakso, Päivi Pajukanta, Jian Yang, John Witte, Noah Zaitlen

## Abstract

Large-scale cohorts with combined genetic and phenotypic data, coupled with methodological advances, have produced increasingly accurate genetic predictors of complex human phenotypes called polygenic risk scores (PRS). In addition to the potential translational impacts of identifying at-risk individuals, PRS are being utilized for a growing list of scientific applications including causal inference, identifying pleiotropy and genetic correlation, and powerful gene-based and mixed model association tests. Existing PRS approaches rely on external large-scale genetic cohorts that have also measured the phenotype of interest. They further require matching on ancestry and genotyping platform or imputation quality. In this work we present a novel reference-free method to produce PRS that does not rely on an external cohort. We show that naive implementations of reference-free PRS either result in substantial over-fitting or prohibitive increases in computational time. We show that our algorithm avoids both of these issues, and can produce informative in-sample PRS over any existing cohort without over-fitting. We then demonstrate several novel applications of reference-free PRS including detection of pleiotropy across 246 metabolic traits and efficient mixed-model association testing.

## INTRODUCTION

Individual genetic polymorphisms typically explain only a small proportion of the heritability, even for traits that are highly heritable [Nolte et al. (2017)]. Polygenic risk scores (PRS), aggregate the contributions of multiple genetic variants to a phenotype [Torkamani et al. (2018)]. These scores can be calculated using routinely recorded genotypes [Torkamani et al. (2018); Nolte et al. (2017)], are strongly associated with heritable traits [Nolte et al. (2017)], and are independent of environmental exposures or other factors that are uncorrelated with germ line genetic variants. These properties have motivated a rapidly expanding list of applications from basic science (e.g. causal inference and Mendelian randomization [Burgess and Thompson (2013)], hierarchical disease models [Cortes et al. (2017)], and identification of pleiotropy [Krapohl et al. (2016)]) to translation (e.g. estimating disease risk [Khera et al. (2018); Maas et al. (2016)], identifying patients who are likely to respond well to a particular therapy [Natarajan et al. (2017)], or flagging subjects for modified screening [Seibert et al. (2018)]).

Polygenic risk scores are calculated as a weighted sum of genotypes. In some applications all genotyped SNPs may be used, but often only a small set are given nonzero weight. A subset of SNPs selected to contribute to a PRS may be a genome-spanning-but-uncorrelated (LD-pruned) set or a set of SNPs with independent evidence of association with the phenotype of interest. Gene-specific polygenic risk scores are also generated using selected sets of SNPs within a region of the genome, such as a window around the coding region of a particular gene [Gusev et al. (2016); Gamazon et al. (2015)]. The weights on the SNPs included in a polygenic score are often derived from the marginal regression coefficients of an external GWAS [Dudbridge (2016); Wray et al. (2007)], but they may instead be based on predictive models using all SNPs. Joint predictive models include LMMs and their sparse extensions [Yang et al. (2011); Zhou et al. (2013); Vilhjálmsson et al. (2015)] and other regularized regression models such as the lasso or elastic net [Warren et al. (2014); Gusev et al. (2016); Gamazon et al. (2015); Rakitsch et al. (2012)]. The predictions from these joint analyses using genome wide variation are also approximated by post-processing of GWAS summary statistics [Vilhjálmsson et al. (2015); Gusev et al. (2016)].

For these SNP weights to accurately reflect the SNPs’ joint association with the phenotype and to generate informative and interpretable polygenic risk scores, the reference data set must match the target data set in many ways: the populations must have similar ancestry; the trait of interest must be measured and in a similar way; and identical genotypes must be assayed or imputed. Further, the reference data must be large enough to accurately learn the PRS weights.

An alternative approach is to use the studied data set to build a reference-free PS. This eliminates the need for an external reference data set with matched genotypes, phenotypes, and populations. However, as we show below, naive approaches can easily over-fit genetic effects. This over-fitting results in PRS correlated with non-genetic components of phenotype, that will induce bias or other errors in downstream applications. Cross-validation is one established approach to mitigate over-fitting, which in this context involves holding out and computing a polygenic score for each sample in turn. The main hurdle to this approach is computation time, as standard leave-one-out cross validation requires fitting the PRS model *N* times in a sample with *N* individuals.

Here we report an efficient method to generate PRS by using the out-of-sample predictions from a cross-validated linear mixed model (LMM). Our approach generates leave-one-out (LOO) polygenic risk scores, which we call *cvBLUPs*, with computational complexity linear in sample size after a single LMM fit. In addition to eliminating the reliance on external data and guaranteeing the PRS are generated from a relevant population and phenotype, we describe several applications that are only feasible with cvBLUPs. We first demonstrate several desirable statistical properties of cvBLUPs and then consider applications including evidence of polygenicity across metabolic phenotypes, a novel formulation of mixed model association studies, and selection of relevant principal components for control of confounding by population structure. To facilitate their use, we have incorporated the calculation of cvBLUPs in the in the genetic analysis program GCTA [Yang et al. (2011)].

## METHODS

We consider the continuous phenotype *y* measured on *N* individuals which depends on an *N*-by-*M* matrix of additively coded genotypes *G*, other covariates *X*, and random noise *ε*:

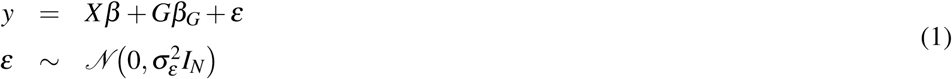

For each subject *i*, the polygenic score PS_*i*_ is calculated as in Equation 2:

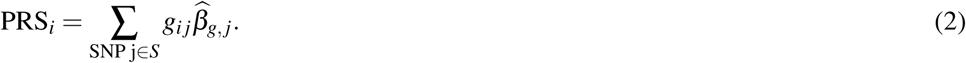

where *S* is the set of SNPs in the polygenic model, *g*_*ij*_ is the number of alleles corresponding to the SNP weights at SNP_*j*_ carried by subject *i*, and 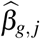 is the SNP weight. 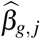 is often chosen to be the estimated effect size of SNP *j* in an external GWAS.

Our objective is to produce a leave-one-out cross-validated polygenic score (PRS) for each subject. We generate our PRS as a genetic prediction from a linear mixed model (LMM). LMMs are widely used for genetic prediction [Robinson et al. (1991)], heritability estimation [Yang et al. (2010); Kang et al. (2008, 2010); Lippert et al. (2011); Zhou and Stephens (2012)], and other polygenic analyses [Lee et al. (2012); Zhou and Stephens (2012); Kang et al. (2008); Lippert et al. (2011)]. The LMM (Equation 3) jointly models the contributions of all SNPs *Z* and other covariates *X* to the phenotype *y*. Following others [Yang et al. (2011)], we define *Z* by centering and scaling columns of *G* to have mean 0 and variance 1.

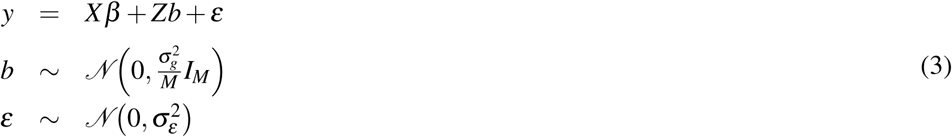

The key LMM parameters are the genetic variance 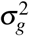 and the noise variance 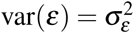. We estimate these by REML [Patterson and Thompson (1971); Kang et al. (2008); Yang et al. (2011)] or Hasman-Elston regression [Chen (2014)]. These variance estimates are then used to estimate *b*, the genetic effect sizes (i.e. weights), or *Zb*, the genetic predictions (i.e. BLUPs):

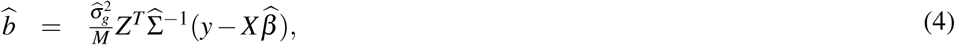

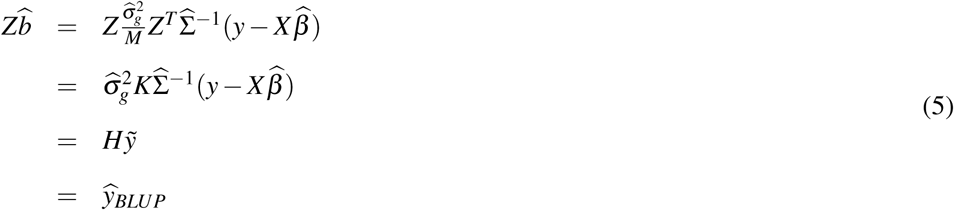

where *K* is the genetic relatedness matrix (GRM) computed from the centered and scaled genotypes *Z* by 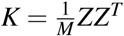. The estimated phenotypic covariance, 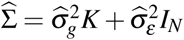, where *I*_*N*_ is the *N*-by-*N* identity matrix, decomposes the covariance to components due to shared genetics (*Zb*) and to noise (*ε*).

While the BLUPs in 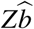 could be used as a polygenic score, we show below that this in general over-fits the noise (*ε*) and is therefore inappropriate for most PRS applications. To address this problem, we propose to use leave-one-out (LOO) cross-validated BLUPs instead of ordinary BLUPs, which guarantees independence between genetic predictions and *ε*. Unfortunately, standard LOO approaches will multiply computational time by a factor of *N*.

We avoid this penalty by leveraging the fact that for linear models, where fitted values are a linear transformation of phenotypes, 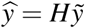, the LOO prediction errors can be calculated from a single model fit [Hastie et al. (2009)]. BLUPs from LMMs fall in this category of linear predictors by applying 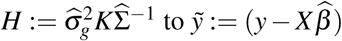, i.e. the phenotype after removing estimated fixed effects.

In more detail, the LOO prediction errors are the differences between the LOO genetic predictions and the observed residual phenotypes after subtracting fixed effects, 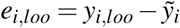. The residuals *r* are the difference between the BLUPs and the residual phenotypes 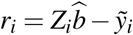. For a linear model, these are related by a simple equation [Hastie et al. (2009)]:

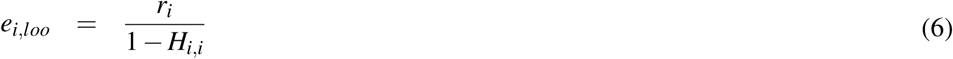

where *H*_*i,i*_ is the *i*’th diagonal element of the matrix *H*. Intuitively this says that due to over-fitting, the in-sample residuals *r*_*i*_ are deflated by (1 − *H*_*i,i*_) relatively to their unbiased LOO counterparts.

We can rearrange these expressions to calculate the LOO predictions, or cvBLUPs, given the standard BLUPs, the phenotype residuals 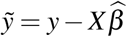, and the diagonal elements of the *H* matrix:

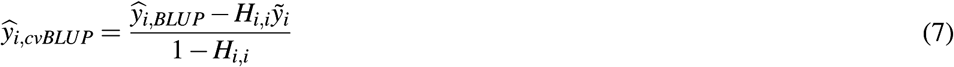

Because all of these elements are computed when fitting an LMM, cvBLUPs can be produced with no additional computational complexity.

## RESULTS

### Empirical confirmation of cross-validated predictions

To examine the properties of the proposed cvBLUP formulation we conducted a set of simulations. We generated 1000 data sets with *N* = 1000 subjects under the model *y* = *Xβ* + *gβ*_*g*_ + *Zb* + *ε*. *X* consists of 5 normally distributed covariates and *Xβ* jointly explain 20% of the phenotypic variance. *g* represents an additively coded SNP with allele frequency 0.5, and *gβ*_*g*_ contributes 2% of the phenotypic variance. *Z* represents *M* = 1000 independent SNPs with minor allele frequencies drawn i.i.d. and uniformly from [0.05, 0.5]. Effect sizes *b*_*j*_ are drawn i.i.d. from 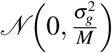 with the genetic variance 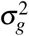 accounting for 39% of the phenotypic variance. The residual noise 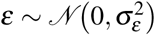 also accounts for 39% of the phenotypic variance, giving a heritability *h*^2^ ≈ 50%. For each simulated data set, we first estimate variance components and then compute BLUPs and cvBLUPs as described above.

Figure 1 shows scatter plots for one simulation of non-genetic (*ε*) and genetic factors (*Zb*) plotted against in-sample BLUPs (Left) and cvBLUPs (Right). As expected, the standard BLUPs are highly correlated with *ε* but the cvBLUPs are not, a central required property of genetic predictors for most applications. In contrast, both BLUPs and cvBLUPs are highly correlated with the true *Zb*, emphasizing the subtlety in constructing valid polygenic risk scores.

**Figure 1.**
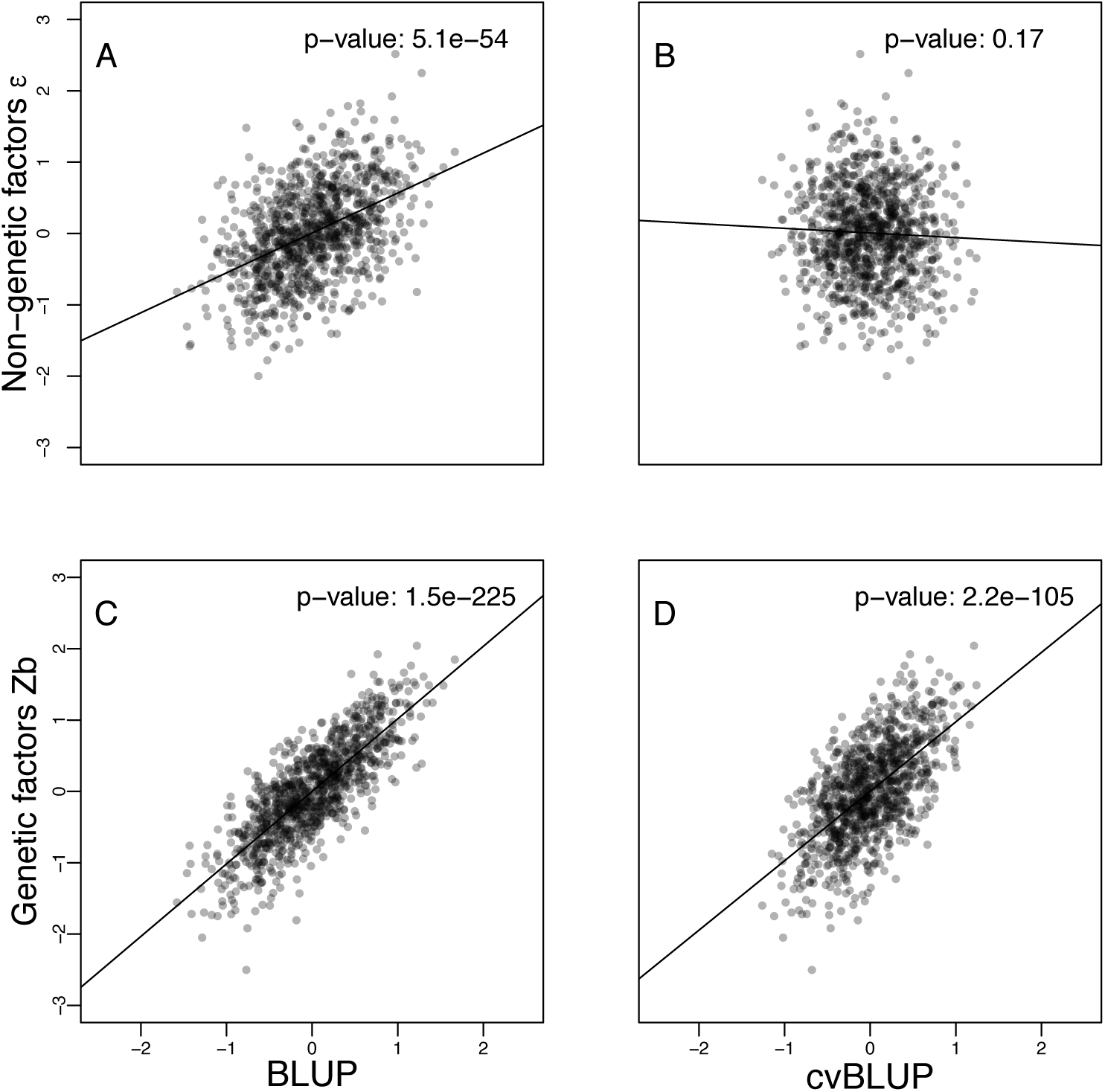
Correlations of genetic predictions, BLUP and cvBLUP, with true genetic factors *Zb* and independent environmental factors *ε* in a simulation of a continuous phenotype with *h*^2^ ≈ 50%, 1000 subjects, and 1000 independent SNPs having random effect sizes. BLUPs are correlated with *ε* while cvBLUPs are not. Lines and p-values are from linear regression fits. *R*^2^ values: A:0.21, B:0.0019, C:0.64, D:0.38.

Table 1 shows the mean correlations of true simulated values with standard BLUPs and cvBLUPs. Again, standard BLUPs are clearly correlated with the noise term *ε* due to over-fitting, but cvBLUPs are appropriately uncorrelated with *ε*. Standard BLUPs, but not cvBLUPs, are also correlated with the unmodeled causal SNP. This type of correlation causes downstream problems for residual analyses, predictions, and causal inference. Importantly, the cvBLUPs are independent of all unmodeled effects as desired.

**Table 1.** Mean correlations (and standard errors) of BLUPs and cvBLUPs with each component of the additive simulation model, *y* = *Xβ* + *gβ*_*g*_ + *Zb* + *ε*. Statistically significant correlations (*α* = 0.05) in bold face.

|  | BLUP | cvBLUP |
| --- | --- | --- |
| $y$ | **0.8241 (0.0013)** | **0.4058 (0.0013)** |
| $X\beta$ | 0.0004 (0.0007) | 0.0005 (0.0009) |
| $Zb$ | **0.7884 (0.0005)** | **0.6212 (0.0008)** |
| $\varepsilon$ | **0.4749 (0.0008)** | 0.0009 (0.0012) |
| Unmodeled Causal SNP | **0.0840 (0.0010)** | 0.0009 (0.0010) |
| Unmodeled Null SNP | 0.0006 (0.0010) | 0.0012 (0.0010) |

## Genetic predictions and cross-trait predictions using cvBLUPs

We next applied cvBLUP in an analysis of Finnish men from the METSIM cohort [Laakso et al. (2017)]. This cohort is comprised of 10197 men aged 45 to 73 at recruitment between 2005 and 2010 in Kuopio, Finland. Blood serum samples were collected from each participant, and 228 metabolites in the samples were quantified by NMR. In addition to the metabolites, biometric traits including height and weight, and epidemiological traits such as diagnoses or family history of diabetes and CHD were recorded for a total of 248 phenotypes. Continuous phenotypes were quantile normalized. All samples were genotyped at 665,478 SNPs on the Illumina OmniExpress chip. After removing subjects with missing rates above 5 percent and SNPs with missing rates above 5 percent, 10070 subjects and 609131 SNPs remain.

We initially consider genetic predictions of the metabolic, biometric, and epidemiological traits in an unrelated subset of subjects (with genetic relatedness less than 0.05). Since the metabolic traits are expected to be affected by statins and by pharmaceutical interventions for diabetes, we exclude subjects with diabetes or who use statins from the initial analysis and calculation of cvBLUPs. There are no comparable data sets with the set of metabolic measurements available in the METSIM cohort, but cvBLUPs allow computationally efficient genetic predictions of all 246 phenotypes (excluding diabetes status and statin use). With the genetic prediction models learned in the restricted set of subjects, we extended predictions to the excluded subjects with standard BLUP effect size estimates (Equation 4). Thus, cvBLUPs allow analyses of reference-free genetic predictions in a subset of subjects that is restricted to avoid confounding by known environmental exposures (statins and responses to diagnoses of diabetes) or by family structure; and these genetic predictions may then be extended to subjects who are initially excluded to avoid confounding.

We first estimated 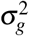 and 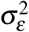 for each phenotype using linear mixed models [Yang et al. (2011)]. Overall, 198 of 246 phenotypes have statistically significant heritability at the 0.05 significance level by Wald tests. The significant heritabilities range from 14.6 to 46.1 percent with a mean and standard deviation of 27.5 and 8.0 percent respectively. The 48 non-significant heritabilities range from 0 to 14.4 percent with a mean and standard deviation of 8.3 and 4.3 percent respectively. We next used the method described above to compute leave-one-out polygenic risk scores (i.e. cvBLUPs). Figure 2 and Supplemental Table 12 show the correlation of the phenotypes (rows) with cvBLUPs (columns) for all 246 phenotypes grouped by hierarchical clustering [Kolde and Kolde (2018)] of rows and applying the same permutation to columns. The blue diagonal shows the expected positive correlation between a cvBLUP and its own phenotype with mean 0.065 and standard deviation 0.037. Focusing on the 198 traits with significant heritabilities the correlations of cvBLUP and phenotype have mean 0.078 and standard deviation 0.027.

**Figure 2.**
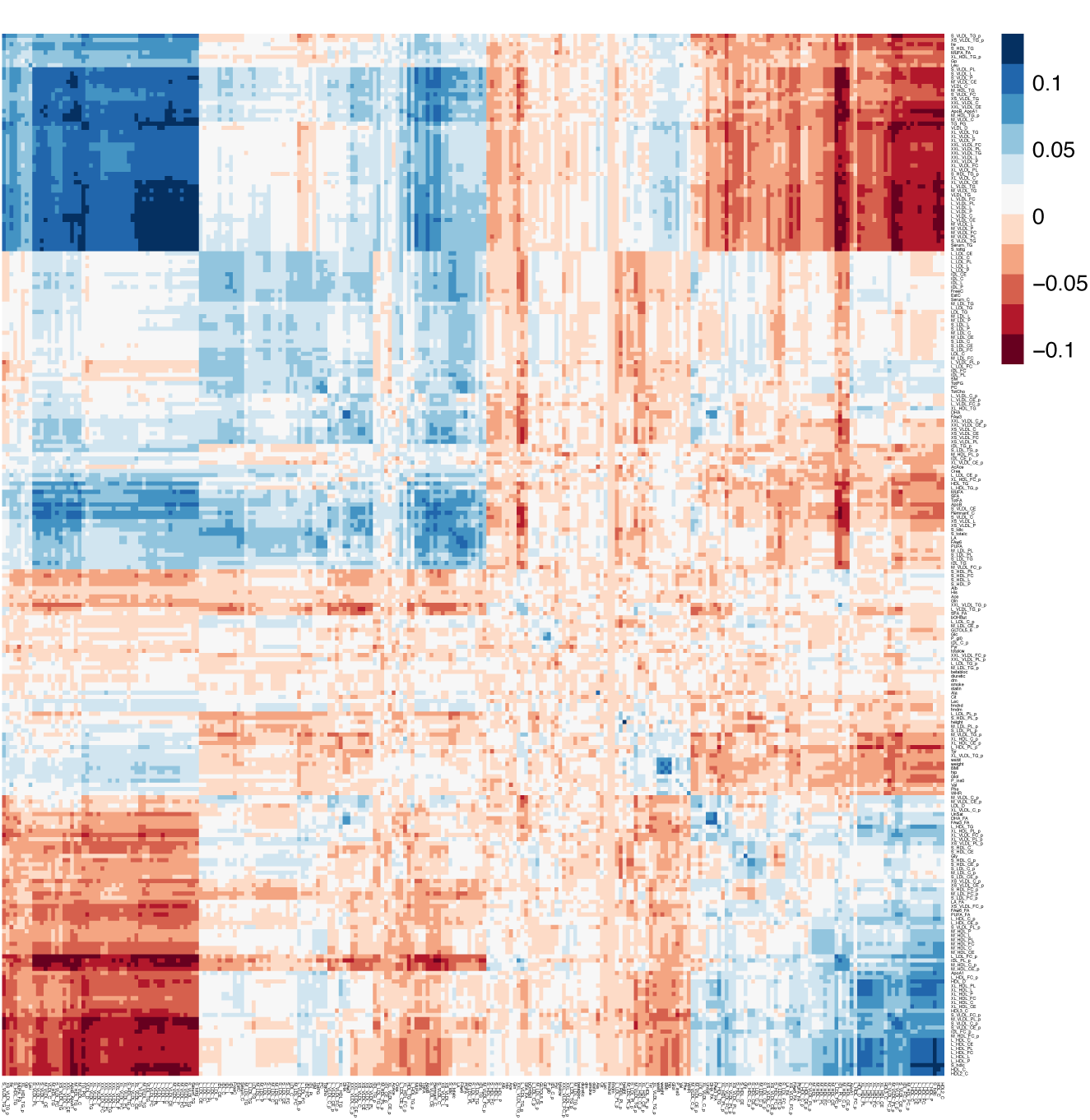
Correlations of phenotypes (rows) and genetic predictions (cvBLUPs, columns) across 246 phenotypes. Many cvBLUPs are strongly correlated with additional phenotypes. A larger version of the figure and a table of the correlations are included in the supplementary materials

The off-diagonal blue patches in the figure represent cvBLUPs that are positively correlated and predictive of different phenotypes, while red patches represent cvBLUPs that are predictive but negatively correlated with different phenotypes. The off-diagonal correlations show the widespread pleiotropy of genetic effects on metabolism with over 16203 off-diagonal cvBLUP-phenotype associations at FDR=0.05 [Benjamini and Yekutieli (2001)]. Many cvBLUP-trait correlations are sign-consistent with the respective trait-trait correlations. For example, HDL and LDL cholesterol are well known be negatively correlated [Terry et al. (1989)], and our results demonstrate this is partially due to negative genetic pleiotropy. That is, we observe that SNPs associated with increased LDL are also associated with decreased HDL in aggregate (and vice versa).

### cvBLUPs in association testing

In addition to use in detection of pleiotropy, polygenic modelling is a widely used tool in association testing [Kang et al. (2008); Yang et al. (2014); Lippert et al. (2011); Zhou and Stephens (2012); Loh et al. (2015)], and we therefore consider cvBLUPs in this this context. We compare the relative performance of five groups of methods. First, unadjusted regression; second, principal component adjusted regression; third, standard linear mixed model (LMM) association tests; fourth, LMM residual based methods; and fifth, cvBLUP adjusted regression. Standard LMM-based methods use association tests where the covariance of the observations based on the genetic relatedness of the subjects is estimated and used to calculate effect estimates and test statistics by generalized least squares [Kang et al. (2008); Yang et al. (2014); Lippert et al. (2011); Zhou and Stephens (2012)]. The LMM residual-based methods perform association tests on the BLUP residualized phenotypes and possibly genotypes. Then, to try and account for the bias inherent in using the overfit BLUPs, they perform an adjustment step on the resulting test statistics and effect sizes. These methods were pioneered by GRAMMAR/GRAMMAR-*γ* and include the recent BOLT and BOLT-INF methods [Aulchenko et al. (2007); Svishcheva et al. (2012); Loh et al. (2015)].

The typical sources of confounding for associations with germ-line genetic markers are population structure, family structure, and batch effects in the data collection [Listgarten et al. (2010)]. Genetic principal components as adjustment covariates may suffice to control for confounding by population structure or batch effects but linear mixed models are often more effective at controlling these sources of confounding [Kang et al. (2010)], while also helping to control confounding by family structure and boosting power to detect true associations over standard fixed-effect regression models [Yang et al. (2014)]. These benefits of linear mixed models come at the expense of an increased computational burden over standard linear regression.

In genome wide association studies the statistical significance of each variant *g*_*j*_ is tested individually. Here, the SNP is jointly analyzed with covariates *X* and the contributions of unmodeled variants contribute to a larger error term *η* = *Z*^(*−j*)^*b*^(*−j*)^ + *ε*.

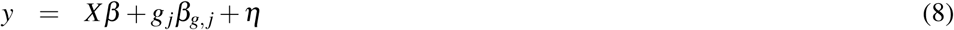

Often a better estimate of the effect size for a particular SNP *g*_*j*_ may be made by accounting for the contributions of the other variants to the phenotype *y*, and by blocking the effects of confounders of the associations of genotypes and the phenotype – by adjustment with an appropriate set of fixed effect covariates or other means [Zaitlen et al. (2012a,b); Yang et al. (2014)].

Here we demonstrate the use of cvBLUPs as adjustment covariates in a linear regression model that efficiently captures some of the benefits of a standard mixed model association study. To compare the performance cvBLUP adjusted analyses to existing methods for association testing under a range of study scenarios, simulations were used. Methods compared were: unadjusted linear regression, PC adjusted linear regression, a standard LMM association test (GCTA), and BOLT association tests. BOLT results were collected both for the infinetissimal genetic model (BOLT-INF) and the sparse causal genetic model (BOLT-LMM). Association tests conducted with cvBLUP adjustment, GCTA, and BOLT were done with leave-one-chromosome-out schemes wherein the variance components, cvBLUPs, and phenotypic predictions and residuals (BOLT) were calculated using only SNPs that are on different chromosomes than the test-SNPs.

In each simulation, data sets were generated with *N* = 2000 subjects under the model *y* = *Xβ* + *gβ*_*g*_ + *Zb* + *ε*. Here *X* consists of normally distributed covariates drawn to contribute i.i.d. noise to the phenotypes in the independent-subject simulation, but to be correlated with the subjects’ ancestral populations in the simulations with population structure and with family in the simulations with confounding by family structure. *Xβ* was scaled to contribute 10% of the phenotypic variance. *g* represents a set of 5 additively coded causal SNPs with effect sizes *β*_*g*_ set to a common fixed value of 0.25. A set of 5 null SNPs were also drawn but did not contribute to the phenotype *y*. *Z* represents *M* = 2000 independent SNPs modeled with random effect sizes. For simulations run under the infinitessimal genetic model, the random effects *b*_*j*_ were drawn from 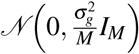 with the genetic variance 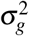 accounting for 40% of the phenotypic variance. For the sparse non-infinitessimal model, a fraction *m*_*c*_ = 2% of the SNPs in *Z* were selected to be causal with effect sizes drawn from 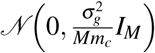 The residual noise 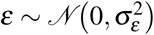 accounts for 40% of the phenotypic variance, giving a heritability *h*^2^ ≈ 50%. Effect estimates and test statistics produced by the various analysis methods are summarized for 1000 null SNPs and 1000 causal SNPs.

In Table 2, the results of analyzing data sets with independent subjects and an infinitessimal genetic architecture are shown. All methods produced unbiased effect estimates and well calibrated tests under the null 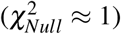, but the cvBLUP and mixed-model based methods were more powerful – with greater average test statistics for the causal SNPs. The BOLT effect estimates were biased, with effect sizes deflated towards zero. This deflation of BOLT effect estimates is seen across the simulation scenarios, however we do not detect deflation of the BOLT effect estimates in the real-data analyses of the METSIM cohort data below, where there are much ratios of counts of SNPs to number of subjects. Bias in BOLT may be due to the empirical estimation of a deflation-correction factor for the residual based test based on the GRAMMAR-*γ* adjusted residual analysis method Loh et al. (2015); Svishcheva et al. (2012). In analogous simulations with a sparse genetic model (Table 9, Supplement), BOLT-LMM is considerably more powerful because it explicitly models the sparse genetic architecture. When the true genetic contribution to the phenotype, *Zb*, (not including the test-SNPs) is included as a covariate in a linear regression association test of causal SNPs *g*, the power is greatly increased.

**Table 2.** Simulations with infinitesimal genetic model, without population or family structure. Mean and standard errors for effect estimates and test statistics of association tests at 1000 Null SNPs with true 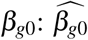 and 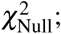; and at 1000 causal SNPs with an alternative hypothesis of true 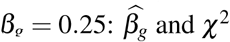 and *χ*^2^

| Model | mean($\beta_{g0}$) | se($\beta_{g0}$) | mean($\chi^2_{Null}$) | se($\chi^2_{Null}$) | mean($\beta_g$) | se($\beta_g$) | mean($\chi^2$) | se($\chi^2$) |
| --- | --- | --- | --- | --- | --- | --- | --- | --- |
| LR | -4.00E-04 | 1.30E-03 | 0.99 | 0.05 | 0.250 | 1.30E-03 | 47.11 | 0.66 |
| LR + 4 PCs | -5.00E-04 | 1.30E-03 | 0.99 | 0.05 | 0.250 | 1.30E-03 | 46.97 | 0.66 |
| LR + cvBLUP | -1.30E-03 | 1.10E-03 | 1.00 | 0.05 | 0.249 | 1.20E-03 | 58.53 | 0.81 |
| LR + BLUP | -1.00E-03 | 6.00E-04 | 1.05 | 0.05 | 0.096 | 6.00E-04 | 36.63 | 0.56 |
| LMM | -1.30E-03 | 1.10E-03 | 1.00 | 0.05 | 0.249 | 1.20E-03 | 56.62 | 0.76 |
| BOLT-INF | -1.30E-03 | 1.00E-03 | 1.01 | 0.05 | 0.219 | 1.00E-03 | 56.13 | 0.76 |
| BOLT-LMM | -1.30E-03 | 1.00E-03 | 1.00 | 0.05 | 0.219 | 1.00E-03 | 56.04 | 0.75 |
| LR + true genetic effect | -1.00E-03 | 9.00E-04 | 1.04 | 0.05 | 0.250 | 9.00E-04 | 96.28 | 1.26 |

In Table 3, the results of analyzing data sets with confounding by population structure are shown. Subjects were drawn from 5 distinct populations with pairwise Fst set to 0.03. Population specific allele frequencies were generated by the Balding-Nickels model [Balding and Nichols (1995)]. The component *Xβ* in the generating model was drawn to be correlated with population, but *X* was not used in the analyses. In this scenario, unadjusted linear regression and linear regression with fewer than 4 principal components as adjustment covariates give inflated test statistics and excessive false positives under the null, 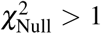. Correcting for population structure by inclusion of 4 principal components, cvBLUP adjustment, or use of the mixed model based methods gives well calibrated test statistics under the null, 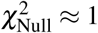. For causal test SNPs, cvBLUP and mixed model based methods give greater test statistics than PC-adjusted linear regression with 4 PCs, indicating greater power. Linear regression with adjustment with the true genetic effect *Zb* gives high power for detecting causal SNPs, but does not control for inflation of test statistics at null SNPs due to population structure 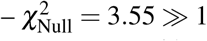. Covariate adjustment with cvBLUPs controls the confounding by population structure and improves the power as do the LMM based methods. These methods use all SNPs and detect the shifts in allele frequencies across populations when there is confounding by population structure.

**Table 3.** Simulations with infinitesimal genetic model and population structure. Mean and standard errors for effect estimates and test statistics of association tests at 1000 Null SNPs with true 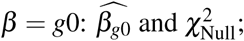 and 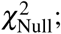; and at 1000 causal SNPs with an alternative hypothesis of true *β*_*g*_ = 0.25: 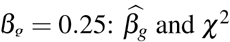 and *χ*^2^

| Model | mean($\beta_{g0}$) | se($\beta_{g0}$) | mean($\chi^2_{Null}$) | se($\chi^2_{Null}$) | mean($\beta_g$) | se($\beta_g$) | mean($\chi^2$) | se($\chi^2$) |
| --- | --- | --- | --- | --- | --- | --- | --- | --- |
| LR | -1.2E-03 | 2.30E-03 | 2.94 | 0.14 | 0.250 | 2.20E-03 | 47.28 | 0.92 |
| LR + 1 PC | -8.00E-04 | 2.10E-03 | 2.44 | 0.12 | 0.250 | 1.90E-03 | 48.20 | 0.85 |
| LR + 2 PCs | -1.00E-04 | 1.80E-03 | 1.94 | 0.09 | 0.253 | 1.70E-03 | 49.00 | 0.80 |
| LR + 3 PCs | -3.00E-04 | 1.60E-03 | 1.49 | 0.07 | 0.252 | 1.50E-03 | 48.90 | 0.76 |
| LR + 4 PCs | 1.00E-03 | 1.30E-03 | 1.02 | 0.05 | 0.250 | 1.30E-03 | 47.86 | 0.70 |
| LR + cvBLUP | 4.00E-04 | 1.20E-03 | 1.00 | 0.04 | 0.240 | 1.20E-03 | 56.8 | 0.81 |
| LMM | -2.00E-04 | 1.20E-03 | 0.99 | 0.05 | 0.252 | 1.20E-03 | 56.97 | 0.79 |
| BOLT-INF | -1.00E-04 | 1.00E-03 | 0.97 | 0.04 | 0.214 | 1.00E-03 | 54.34 | 0.76 |
| BOLT-LMM | -1.00E-04 | 1.00E-03 | 0.97 | 0.04 | 0.214 | 1.00E-03 | 54.37 | 0.76 |
| LR + true genetic effect | 2.20E-03 | 1.80E-03 | 3.55 | 0.16 | 0.250 | 1.80E-03 | 89.28 | 1.56 |

cvBLUPs correct for population structure because they are weighted combinations of ALL principal components, where weights are based on the the singular value corresponding to the principal component and on the strength of the association of the principal component with the outcome (see Supplement). Conceptually, cvBLUPs control for population structure as if all PCs were considered and the most relevant ones for the analysis were kept.

In Table 4, the results of analyzing data sets with confounding by family structure are shown. Here the 2000 subjects in each simulation represented 200 families with 10 subjects each. Families were generated in pedigrees with four founders and six of their descendants, with descendants’ genotypes selected independently by drop-down from their parents. In the data generating model, there were covariate effects correlated with family membership, *Xβ* but these covariates were not included in the analyses, creating confounding by family structure. In this scenario, unadjusted linear regression, and PC-adjusted linear regression have inflated test statistics 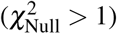 and correspondingly high false discovery rates. Standard linear mixed models (GCTA) control for the confounding by family structure, with accurate effect estimates 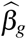 under the null and alternate, and barely inflated test statistics under the null 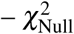 having mean and standard deviation 1.10 and 0.05 respectively. In this scenario, cvBLUP adjusted analyses and the results from BOLT have biased effect estimates under the alternative, and deflated (conservative) test statistics under the null. This suggests over-adjustment for family structure in both cvBLUP-adjusted analyses and BOLT.

**Table 4.** Simulations with infinitessimal model and family structure. Mean and standard errors for effect estimates and test statistics of association tests at 1000 Null SNPs with true 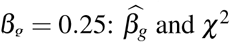 and *χ*^2^

| Model | mean($\beta_{g0}$) | se($\beta_{g0}$) | mean($\chi^2_{Null}$) | se($\chi^2_{Null}$) | mean($\beta_g$) | se($\beta_g$) | mean($\chi^2$) | se($\chi^2$) |
| --- | --- | --- | --- | --- | --- | --- | --- | --- |
| LR | -1.30E-03 | 1.60E-03 | 1.52 | 0.07 | 0.252 | 1.60E-03 | 48.81 | 0.77 |
| LR + 4 PCs | -5.00E-04 | 1.50E-03 | 1.49 | 0.06 | 0.253 | 1.60E-03 | 48.93 | 0.75 |
| LR + cvBLUP | 4.00E-04 | 1.00E-03 | 0.81 | 0.03 | 0.183 | 9.00E-04 | 36.20 | 0.52 |
| LMM | 5.00E-04 | 1.40E-03 | 1.10 | 0.05 | 0.251 | 1.30E-03 | 48.57 | 0.68 |
| BOLT-INF | 5.00E-04 | 9.00E-04 | 0.77 | 0.03 | 0.166 | 9.00E-04 | 33.65 | 0.46 |
| BOLT-LMM | 5.00E-04 | 9.00E-04 | 0.84 | 0.04 | 0.166 | 9.00E-04 | 36.24 | 0.51 |
| LR + true genetic effect | 5.00E-04 | 1.00E-03 | 1.21 | 0.05 | 0.252 | 1.00E-03 | 98.24 | 1.31 |

Finally, we applied these methods to the METSIM data described above. All p-values were GC adjusted for comparison purposes. All mixed model-based methods, including LR+cvBLUP were more powerful than standard linear regression. As expected BOLT-LMM had the highest power due to modelling of non-infinitessimal structure. In this data analysis there is no observed evidence of bias in effect size estimates by BOLT, such as a systematic deflation of effect estimates relative to estimates made by standard linear regression or standard LMM fixed effect estimates, suggesting that it may not be a problem in practice.

**Table 5.**
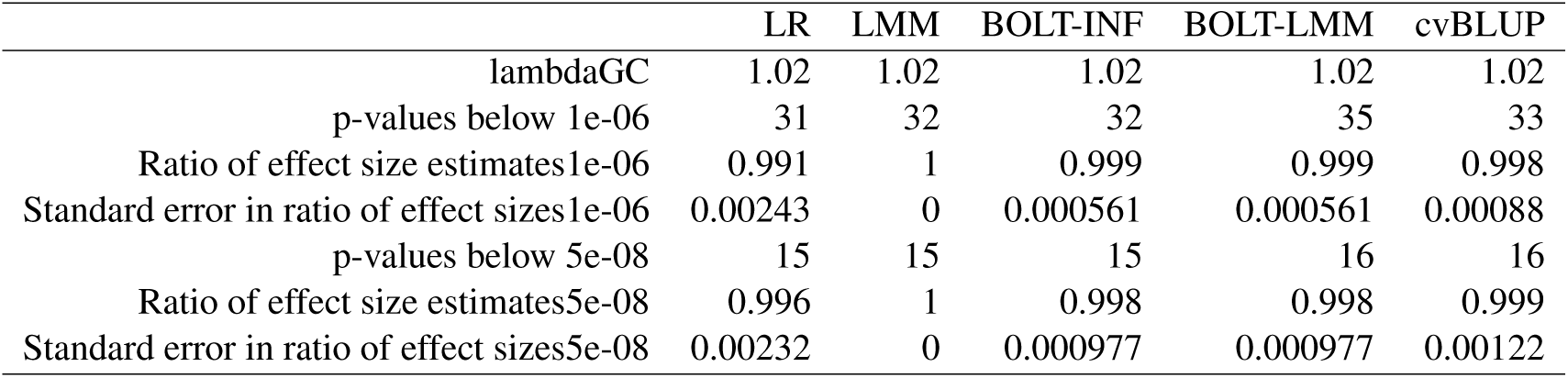
GWAS results for baseline low-density lipoprotein cholesterol (LDLC) using an unrelated subset of subjects from the METSIM cohort. LR: linear regression, LMM: linear mixed model with GLS analysis of SNP effects implented in GCTA, cvBLUP: cross-validated prediction-adjusted linear regression, BOLT-INF; BOLT assuming infinitesimal genetic model, BOLT-LMM: BOLT using mixture of Normal distributions as prior for SNP effect sizes, i.e. sparse genetic architecture. cvBLUP-adjusted analyses, LMM, and BOLT were used in a leave-one-chromosome-out scheme with variance components, cvBLUPs, covariance models (LMM, GCTA), and genetic predictions and residuals (BOLT) generated using SNPs on chromosomes other than that of the test-SNPs.

|  | LR | LMM | BOLT-INF | BOLT-LMM | cvBLUP |
| --- | --- | --- | --- | --- | --- |
| lambdaGC | 1.02 | 1.02 | 1.02 | 1.02 | 1.02 |
| p-values below 1e-06 | 31 | 32 | 32 | 35 | 33 |
| Ratio of effect size estimates1e-06 | 0.991 | 1 | 0.999 | 0.999 | 0.998 |
| Standard error in ratio of effect sizes1e-06 | 0.00243 | 0 | 0.000561 | 0.000561 | 0.00088 |
| p-values below 5e-08 | 15 | 15 | 15 | 16 | 16 |
| Ratio of effect size estimates5e-08 | 0.996 | 1 | 0.998 | 0.998 | 0.999 |
| Standard error in ratio of effect sizes5e-08 | 0.00232 | 0 | 0.000977 | 0.000977 | 0.00122 |

**Table 6.**
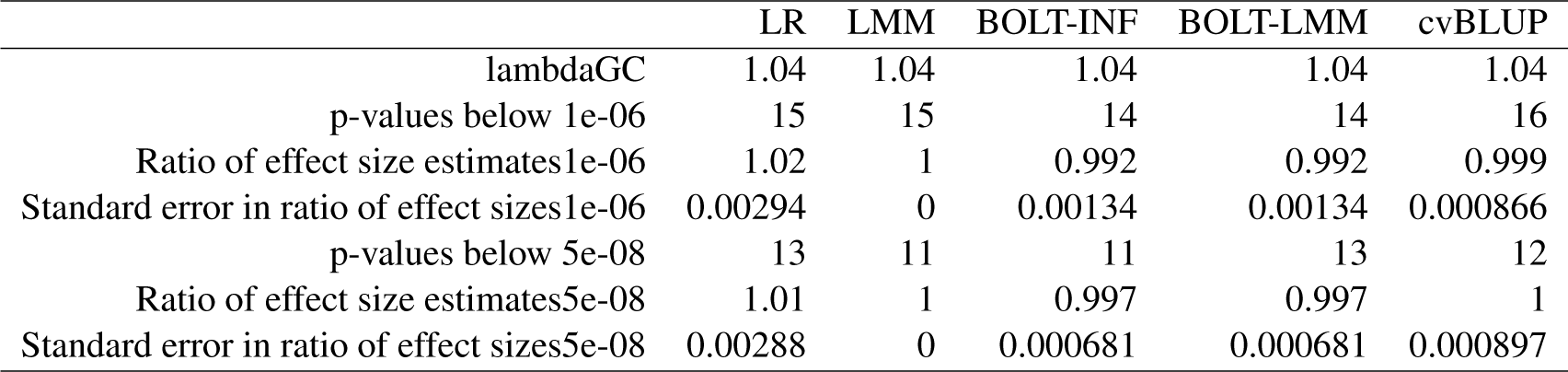
GWAS results for baseline high-density lipoprotein cholesterol (HDLC) using an unrelated subset of subjects from the METSIM cohort. LR: linear regression, LMM: linear mixed model with GLS analysis of SNP effects implemented in GCTA, cvBLUP: cross-validated prediction-adjusted linear regression, BOLT-INF; BOLT assuming infinitesimal genetic model, BOLT-LMM: BOLT using sparse genetic architecture.cvBLUP-adjusted analyses, LMM, and BOLT were used in a leave-one-chromosome-out scheme with variance components, cvBLUPs, covariance models (LMM, GCTA), and genetic predictions and residuals (BOLT) generated using SNPs on chromosomes other than that of the test-SNPs.

### Leveraging cvBLUPs to compute linear mixed model shrink parameters

Finally, we examine the use of cvBLUPs in estimating LMM shrink parameters. The BLUP effect size estimates from a linear mixed model are “shrunk” to be smaller in magnitude to exploit a bias-variance trade-off that reduces their mean squared error. With the estimated genetic variance component 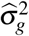, the estimated distribution of effect estimates 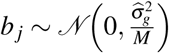 is an empirical prior on the effect sizes, which yields estimates that are biased toward the prior mean of 0. Equivalently, the BLUP effect sizes estimates are ridge regression estimates with a penalty parameter λ related to the LMM variance components: 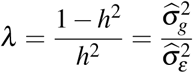.

When SNPs are independent, the LMM shrinkage estimates of the effect sizes *b* for given estimates of the variance components is found to be [Vilhjálmsson et al. (2015)]

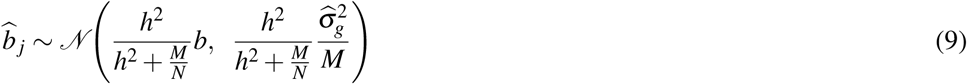

and the shrink, 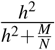, can be calculated directly from the number of individuals and SNPs. However, when SNPs are in LD, it is more complicated to estimate the effective number of SNPs [Patterson et al. (2006)] and hence the shrink. Here we show how cvBLUPs can be used to estimate the shrink directly.

First, in the case of independent SNPs, the LMM estimates of the SNP effects 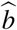 would be reduced in magnitude and variance relative to the true values *b*, as in Equation 9, and there would be a corresponding decrease in the variance of polygenic risk scores for a new individual with normalized genotypes *Z*_new_, calculated using 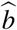 as weights rather than the true effect sizes *b*:

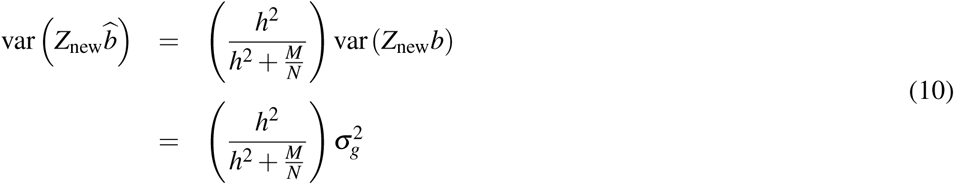

Equation 10 suggests the direct estimation of the shrink by taking the ratio of 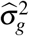 to 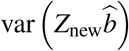. We confirm this approach is approximately unbiased via simulation. For multiple settings of *N* (number of subjects), *M* (number of SNPs) and *h*^2^ (heritability), heritability and variance components were estimated and the value of the standard independent-SNP model for the shrink is compared to the empirical BLUP shrink 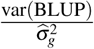 and the empirical cvBLUP shrink 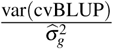. In each simulation setting, the BLUP shrink is much larger than the independent model value or the cvBLUP shrink due to over-fitting by the standard BLUPs. However, the cvBLUP formula and the independent-SNP model are consistent for all parameter settings.

Because this approach does not require identification of an effective number of SNPs, it extends directly to the case where there is linkage disequilibrium. We applied this approach to simulated data from the METSIM cohort used above. We estimated the shrink parameter for simulated phenotypes based on the real genotypes at 609131 SNPs with minor allele frequencies greater than 0.01 on the 6263 unrelated (at the 0.05 level) subjects. Twenty simulations were run using fractions of causal SNPs between 0.0001 and 1.0. Causal SNPs were chosen by simple random sampling with equal probability from the genotyped SNPs. SNP effect sizes were normally distributed and trait heritability was 50 percent. The shrink formula for independent SNPs suggested a shrink of about 0.0045-0.0055, while the cvBLUP shrink – the ratio of the variance of the cvBLUPs to the estimated genetic variance component ranged from 0.06-0.08.

In this setting, with LD between the SNPs, the independent SNP formula for the shrink is invalid. In particular the effective number of SNPs 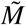 is much smaller than the total number of SNPs leading to a shrink estimate over ten times smaller than the the cvBLUP based estimate.

**Table 7.** Estimation of the shrink parameters for BLUPs and cvBLUPs in simulations with independent SNPs. The independent shrink is that derived in Formula 9 for independent SNPs.

| N | M | $h^2$ | $\hat{h}^2$ | Independent Shrink | BLUP Shrink | cvBLUP Shrink |
| --- | --- | --- | --- | --- | --- | --- |
| 400 | 400 | 0.5 | 0.512 (0.102) | 0.335 (0.003) | 0.627 (0.007) | 0.389 (0.006) |
| 400 | 800 | 0.5 | 0.502 (0.119) | 0.200 (0.003) | 0.560 (0.010) | 0.217 (0.005) |
| 400 | 1200 | 0.5 | 0.494 (0.128) | 0.139 (0.003) | 0.530 (0.011) | 0.150 (0.004) |
| 800 | 400 | 0.5 | 0.492 (0.062) | 0.496 (0.002) | 0.695 (0.004) | 0.573 (0.004) |
| 800 | 800 | 0.5 | 0.501 (0.066) | 0.334 (0.002) | 0.619 (0.005) | 0.383 (0.004) |
| 800 | 1200 | 0.5 | 0.500 (0.075) | 0.248 (0.002) | 0.581 (0.006) | 0.280 (0.004) |
| 1200 | 400 | 0.5 | 0.505 (0.052) | 0.600 (0.002) | 0.770 (0.003) | 0.700 (0.003) |
| 1200 | 800 | 0.5 | 0.501 (0.051) | 0.426 (0.002) | 0.661 (0.003) | 0.494 (0.003) |
| 1200 | 1200 | 0.5 | 0.501 (0.051) | 0.332 (0.002) | 0.614 (0.004) | 0.378 (0.003) |
| 400 | 400 | 0.1 | 0.105 (0.080) | 0.089 (0.006) | 0.199 (0.010) | 0.106 (0.006) |
| 400 | 800 | 0.1 | 0.114 (0.096) | 0.052 (0.004) | 0.174 (0.011) | 0.060 (0.004) |
| 400 | 1200 | 0.1 | 0.114 (0.104) | 0.035 (0.003) | 0.170 (0.011) | 0.044 (0.003) |
| 800 | 400 | 0.1 | 0.102 (0.044) | 0.165 (0.006) | 0.238 (0.008) | 0.169 (0.006) |
| 800 | 800 | 0.1 | 0.101 (0.056) | 0.090 (0.005) | 0.178 (0.008) | 0.094 (0.004) |
| 800 | 1200 | 0.1 | 0.105 (0.066) | 0.063 (0.004) | 0.160 (0.008) | 0.067 (0.004) |
| 1200 | 400 | 0.1 | 0.098 (0.032) | 0.222 (0.006) | 0.283 (0.007) | 0.225 (0.006) |
| 1200 | 800 | 0.1 | 0.101 (0.035) | 0.130 (0.004) | 0.208 (0.006) | 0.132 (0.004) |
| 1200 | 1200 | 0.1 | 0.098 (0.041) | 0.088 (0.003) | 0.170 (0.006) | 0.090 (0.004) |

**Table 8.** Estimates of the shrink parameter for simulated phenotypes based on the real genotypes METSIM cohort and various genetic architectures or fractions of causal SNPs. The independent shrink is that derived in Formula 9 for independent SNPs.

| Fraction Causal SNPs | Independent Shrink | BLUP Shrink | cvBLUP Shrink |
| --- | --- | --- | --- |
| 1.0 | 0.0050 (1.60E-04) | 0.514 (0.015) | 0.076 (0.002) |
| 0.1 | 0.0053 (1.71E-04) | 0.539 (0.016) | 0.079 (0.002) |
| 0.01 | 0.0055 (1.73E-04) | 0.556 (0.016) | 0.080 (0.002) |
| 0.001 | 0.0047 (5.45E-05) | 0.477 (0.005) | 0.067 (0.001) |
| 0.0001 | 0.0040 (1.20E-04) | 0.409 (0.011) | 0.058 (0.001) |

## DISCUSSION

Here we describe a new and computationally efficient approach for generating polygenic risk scores directly from a linear mixed model. We show that the LMM framework allows direct calculation of out-of-sample genetic predictions. Our approach will have immediate utility for the growing list of applications that rely on PS, and we provide examples of several additional application areas, including detection of pleiotropy, powerful association testing, and estimation of polygenic shrink.

The elimination of over-fitting by cvBLUPs relative to BLUPs suggests a solution to the bias problem in residual based methods. Rather than post-hoc correction of residual test statistics as in GRAMMAR-*g* and BOLT, the LMM residuals may be replaced by out-of-sample prediction errors with cross-validated predictors: use (*y* – cvBLUP) instead of standard residuals (*y* – BLUP)

There are several limitations of this approach. First, cvBLUPs are calculated from the standard LMM framework, which corresponds to the infinitessimal genetic model. As we see in in supplementary Tables 9-11, in simulations with sparse genetic architectures, sparse models have considerably higher power. In future work, we intend to use sparse analogues of cvBLUPs generated as cross-validated predictions. Unfortunately, sparse models including the BOLT-LMM model and LASSO are not amenable to the fast leave-one-out cross-validation as in Equation (7).

Another limitation is that when used as adjustment covariates cvBLUPs do not control confounding by family structure or cryptic relatedness as well as standard mixed model association tests. Rather than using all subjects for computation of cvBLUPs, an alternative protocol is to calculate cvBLUPs for an unrelated subset, and also to calculate BLUP effect size estimates 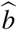 (Equation 4) using these unrelated subjects. This procedure will tend to block confounding by genetic structure remaining in the nominally unrelated subset and will improve power by accounting for the polygenicity of the trait, but it could under-adjust for confounding by family structure because by construction the training set for learning the polygenic model does not contain closely related subjects. Methods involving cross-validated predictions from multiple models or cvBLUPs from mixed models with multiple variance components may prove useful, by analogy with other multiple variance component methods that include sparse relatedness matrices to indicate family membership [Zaitlen et al. (2013), Tucker et al. (2015)]

In our cross-trait analysis of the METSIM data set we show that the cross-correlations of PRS for one trait and actual (normalized) phenotypic measures for other traits. We are working to extend these cross-trait analyses – in particular by using correlations of cvBLUPs for pairs of traits as estimates of the genetic correlation. However, even naive correlations of cvBLUPs give an effective picture of the genetic correlations between traits. Since cvBLUPs are efficiently calculated one at a time, and genetic correlations are then estimated in trivial pairwise analyses of traits, the pair-wise correlations of hundreds or thousands of traits may be efficiently calculated this way. Furthermore, in the pair-wise analyses of the METSIM phenotypes, and in our pair-wise analyses of RNA expression that underlie our trans-eQTL analysis [Liu et al. (2018)] we actually generate asymmetric cross-correlation matrices because the correlation of cvBLUP for trait A with measured trait B is not the same as the correlation of measured trait A with cvBLUP for trait B. We are exploring applications of these asymmetric matrices for network analysis and causal inference.

Efficient generation of out-of-sample genetic predictions using leave-one-out cross-validation of the predictions from a linear mixed model is an effective way to generate polygenic risk scores, and opens the application of analyses based on PRS to scenarios where there is no available reference data to generate a typical scoring model. It is now well known that PRS and genetic predictions transfer poorly to populations that are distinct from the reference data set used to learn the genetic model [Martin et al. (2017); Scutari et al. (2016)]. We look forward to using reference-free PRS methods based on cvBLUPs for applications with data from under-represented populations.

The principle of using cross-validated predictions from polygenic models as PRS may be extended to predictions from sparse or complex models, but the cross-validated predictions from a standard linear mixed model, which we call cvBLUPs, are particularly simple to calculate and have novel and interpretable applications. To make the results of this work accessible to the community, we have implemented them in the GCTA software package [Yang et al. (2011)].

## ACKNOWLEDGEMENTS

NZ, JM, DP were funded by NIH grants (K25HL121295, U01HG009080, R01HG006399).

We thank the individuals who participated in the METSIM study. This study was funded by the National Institutes of Health (NIH) grants HL-095056, HL-28481, and U01 DK105561. The funders had no role in study design, data collection, and analysis, decision to publish, or preparation of the article.

MAK works in a Unit that is supported by the University of Bristol and UK Medical Research Council (MC_U_U_1_2013/1). The Baker Institute is supported in part by the Victorian Government’s Operational Infrastructure Support Program.

## APPENDICES

### Formula for efficient leave-one-out cross-validation of the linear mixed model

Let

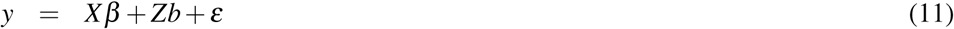

represent a linear mixed model with continuous outcome *y*, fixed effect covariates *X*, fixed effect sizes *b*, additively coded genotypes *Z*, random genetic effect sizes *b* and unmodeled or environmental factors *ε*. The genetic effect sizes *b* and environmental factors *ε* are modeled as i.i.d. normally distributed random variables with variances 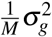 and 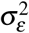 respectively:

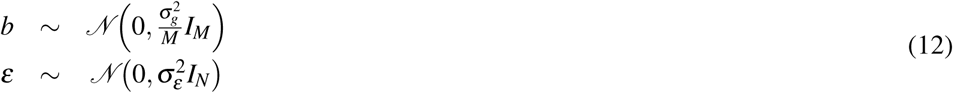

By convention the genotypes are scaled to have mean 0 and variance 1, and the SNP effect sizes are assumed to have effect sizes independent of MAF on this scale. So the total genetic contributions to the phenotype *Zb* are normally distributed with mean 0 and variance 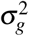. The scaled genotypes *Z* are used to calculate a SNP-based genetic relatedness matrix *K*:

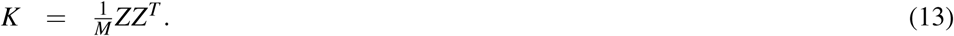

The observations *y* in the linear mixed model are normally distributed with a covariance ∑ that depends on *K* and the variance components 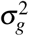 and 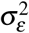

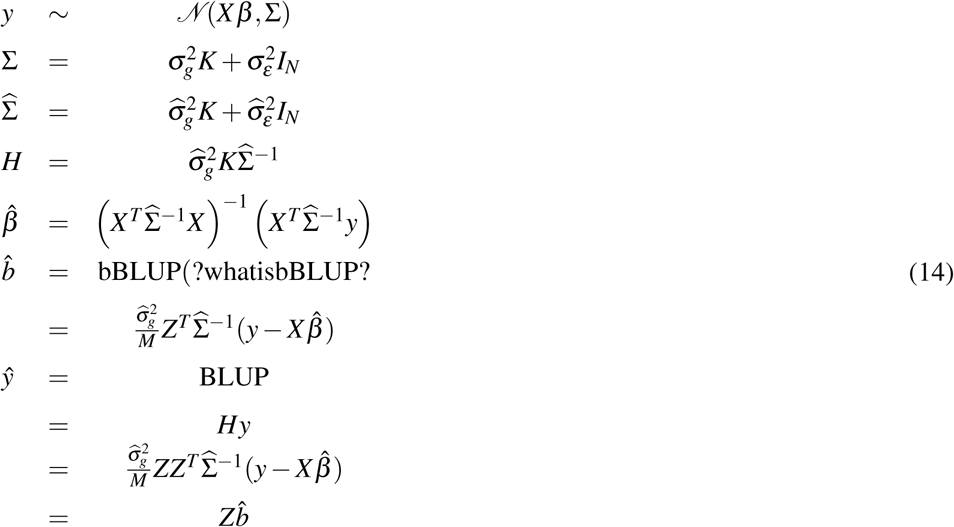

To simplify notation, let fixed effect sizes *β* and fixed effects *Xβ* = 0:

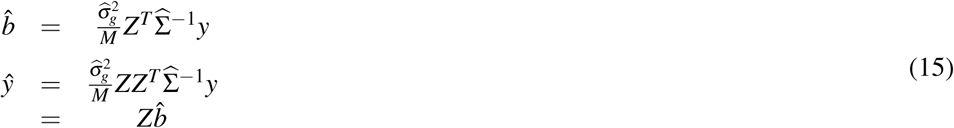

Exclude observation *i* from the genetic predictive model fit and then make an out-of-sample (oos) prediction for observation *i*:

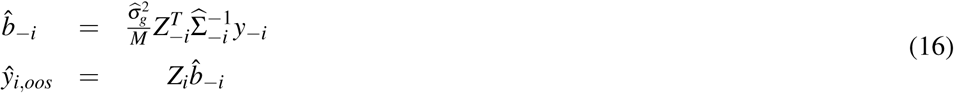

Now generate the augmented vector *y*_+*i*_ as the vector of outcomes *y* with observation *y*_*i*_ replaced with it’s out-of-sample prediction from a model trained on the remaining observations, 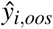.

Consider the ridge regression interpretation of the mixed model with the ridge penalty 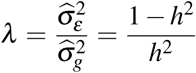. Assume the variance components or heritability have been estimated in a prior step or are known.

For the reduced data set with the outcome for observation *i* removed we have:

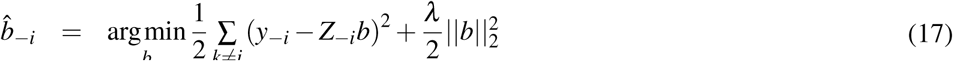

For the augmented data set with the outcome for observation *i* replaced by an out-of-sample prediction we have:

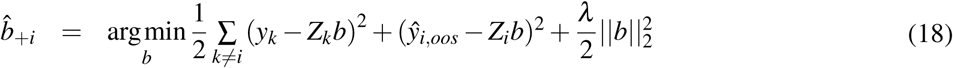

For the augmented model, 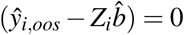, and the remaining terms in the expression for 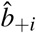 are the same as for the reduced model 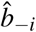, so 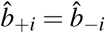.

Now consider the differences in the predictions for the *i*th value from the augmented model and the model with all observations:

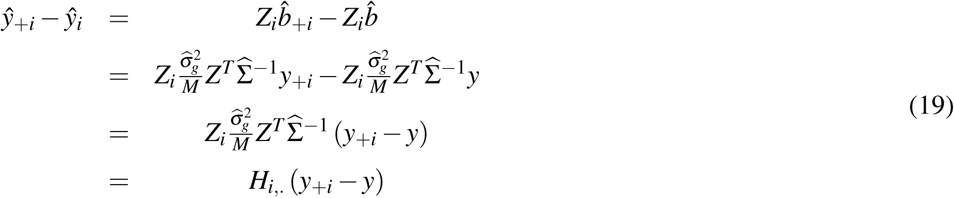

Here, 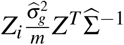 is the *i*th row of the matrix *H*, or *H*_*i*,._. The vectors *y*_+*i*_ and *y* only differ at the *i*th element, so:

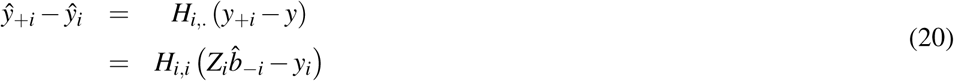

Finally,

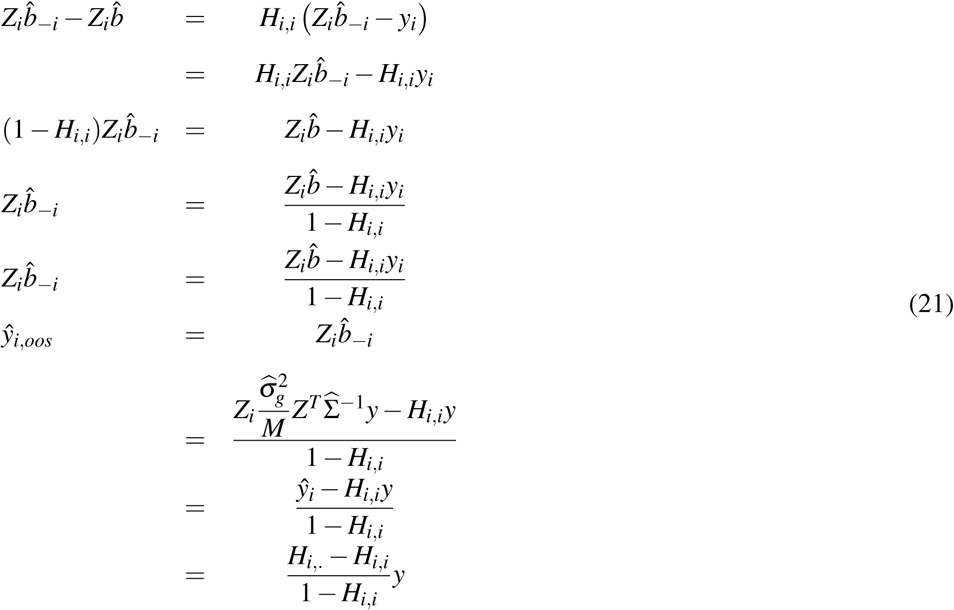

Relationship between principal components and cross-validated BLUPs

Genetic principal components are routinely used as quantitative measures of study participants’ ancestry [Patterson et al. (2006); McVean (2009)] and as such are used as adjustment covariates in association studies to block confounding by population structure [Patterson et al. (2006)]. Linear mixed models provide another framework for controlling potential confounding by population structure [Yang et al. (2014)]. Both PCs and LMMs are methods that account for ancestry and other forms of genetic structure in a data set by analyses of the genetic relatedness matrix (GRM) 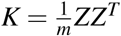. Specifically, PCA involves calculating some number of eigenvetors of the GRM – or equivalently left singular vectors of the scaled genotype matrix *Z* – and using them as adjustment covariates in regression analyses, while LMMs model observed outcomes as non-independent, with the random effects that contribute to the outcome *y* correlated to a degree related to the amount of shared genetic variation between each pair of subjects: 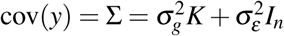.

In principal component adjusted analyses, some number of PCs are used as adjustment covariates in linear regression. With linear mixed models, generalized least squares with an estimate of the sample covariance 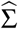 is used.

Principal component adjusted analysis has several disadvantages. First, it is not clear how many principal components should be used. In GWAS it has become conventional to use a standard number of PCs, say 10, but it is generally not clear whether that will be enough to account for the components of genetic structure that are confounded with non-genetic factors in a study. Second, it is not clear which PCs should be included. Conventionally, PCs are sorted in descending order by the magnitude of the corresponding eigenvalues, and PCs with the largest eigenvalues are used. However, selected PCs may not be associated with any non-genetic factors and may not relieve any confounding. Finally, in association testing applications, tests needlessly lose power due to over-adjustment when PCs that do not adjust for confounding are included as covariates. By construction PCs represent axes of variation in the genotype data, so some may be highly correlated with a test SNP, and inclusion of correlated covariates increases the standard errors around test SNP effect estimates.

The spectral decomposition of the GRM *K* is:

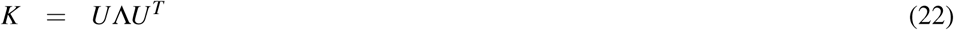

where *U* is a unitary matrix whose columns are the genetic PCs or eigenvectors of *K*, and Λ is a diagonal matrix of the corresponding eigenvalues.

Principal component adjusted regression includes *U*_*k*_ (first *k* columns of *U*) as regression covariates to improve the estimation or testing of *β*_*g*_:

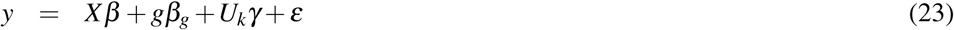

This leaves open the question of how many or which PCs should be included. As an alternative to standard practice, we can try using all of the PCs, and use regularization to keep the model estimable. We can also minimize problems due to over-adjustment by calculating the PCs *U* using a GRM *K\** or scaled genotype matrix *Z\** that does not include SNPs in linkage disequilibrium with or on the same chromosome as the test-SNP *g*.

Towards a connection to cvBLUPs, rescale the columns of *U* by the corresponding singular values:

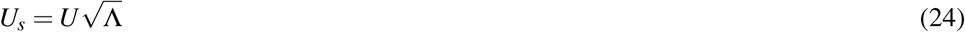

Rescaling *U* to *U*_*s*_ puts a higher prior on PCs with larger eigenvalues. Now compress the contribution of the principal components to the outcome by calculating a vector of phenotypic predictions using ridge regression with a penalty *λ* on the principal components:

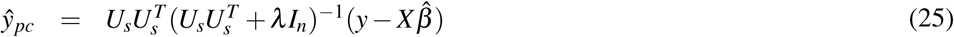

If we define 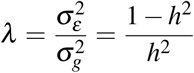 and recall that 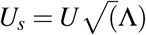, so that 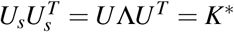, we have:

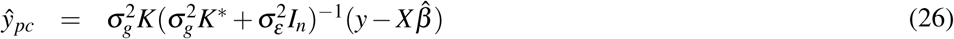

But this is just a BLUP. So, BLUPs arise from a limiting case of trying to do PC-adjusted regression with all PCs. Therefore cross-validated predictions from ridge regression on all PCs are cvBLUPs.

cvBLUPs as adjustment covariates are similar to a compression of all PCs into a single covariate, with the PCs given prior weights that emphasize the PCs with larger eigenvalues, but do not exclude any. The PCs are also weighted by their relevance to the outcome because they represent predictions from a ridge regression model that implicitly has ridge regression effect sizes for the association of the PC with the outcome. Over-fitting in the ridge regression step is avoided by the leave-one-out cross-validation. Finally, loss of power by over adjustment is avoided by excluding the chromosome or SNPs in linkage disequilibrium with test SNP from the GRM used for calculation of the cvBLUPs. In fact, unlike PC adjustment but similar to standard LMM analyses, cvBLUP adjustment boosts the power of association studies by modeling genetic contributions to the phenotype other than the SNP of interest, thereby increasing the signal to noise ratio.

### Additional results with cvBLUP-adjusted association testing

**Table 9.** Simulations with sparse genetic model and independent subjects. Mean and standard errors for effect estimates and test statistics of association tests at 1000 Null SNPs with true 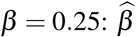 and 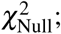; and at 1000 causal SNPs with an alternative hypothesis of true 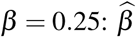 and *χ*^2^

| Model | mean($\beta_0$) | se($\beta_0$) | mean($\chi^2_{Null}$) | se($\chi^2_{Null}$) | mean($\beta$) | se($\beta$) | mean($\chi^2$) | se($\chi^2$) |
| --- | --- | --- | --- | --- | --- | --- | --- | --- |
| LR | -1.10E-03 | 1.40E-03 | 1.14 | 0.05 | 0.249 | 1.30E-03 | 47.48 | 0.70 |
| LR + 4 PCs | -9.00E-04 | 1.40E-03 | 1.13 | 0.05 | 0.249 | 1.30E-03 | 47.38 | 0.68 |
| LR + cvBLUP | -1.10E-03 | 1.30E-03 | 1.14 | 0.05 | 0.248 | 1.20E-03 | 58.15 | 0.83 |
| LMM | -1.10E-03 | 1.30E-03 | 1.14 | 0.05 | 0.248 | 1.20E-03 | 56.22 | 0.78 |
| BOLT-inf | -9.00E-04 | 1.10E-03 | 1.14 | 0.05 | 0.219 | 1.00E-03 | 55.81 | 0.77 |
| BOLT-LMM | -9.00E-04 | 1.10E-03 | 1.09 | 0.05 | 0.219 | 1.00E-03 | 80.99 | 1.05 |
| LR + true genetic effect | -6.00E-04 | 1.00E-03 | 1.13 | 0.05 | 0.249 | 9.00E-04 | 94.32 | 1.26 |

**Table 10.** Simulations with sparse genetic architecture and population structure. Mean and standard errors for effect estimates and test statistics of association tests at 1000 Null SNPs with true 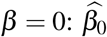 and 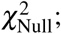; and at 1000 causal SNPs with an alternative hypothesis of true 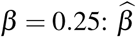 and *χ*^2^

| Model | mean($\beta_0$) | se($\beta_0$) | mean($\chi^2_{Null}$) | se($\chi^2_{Null}$) | mean($\beta$) | se($\beta$) | mean($\chi^2$) | se($\chi^2$) |
| --- | --- | --- | --- | --- | --- | --- | --- | --- |
| LR | -1.50E-03 | 2.30E-03 | 2.88 | 0.14 | 0.245 | 2.20E-03 | 46.30 | 0.90 |
| LR + 4 PCs | -6.00E-04 | 1.30E-03 | 1.01 | 0.05 | 0.248 | 1.20E-03 | 46.57 | 0.66 |
| LR + cvBLUP | -6.00E-04 | 1.10E-03 | 0.97 | 0.05 | 0.238 | 1.10E-03 | 54.76 | 0.78 |
| LMM | -7.00E-04 | 1.20E-03 | 1.00 | 0.05 | 0.247 | 1.20E-03 | 55.22 | 0.76 |
| BOLT-inf | -8.00E-04 | 1.00E-03 | 0.97 | 0.05 | 0.211 | 1.00E-03 | 52.78 | 0.73 |
| BOLT-LMM | -8.00E-04 | 1.00E-03 | 1.02 | 0.05 | 0.211 | 1.00E-03 | 77.75 | 1.05 |
| LR + true genetic effect | -2.30E-03 | 1.90E-03 | 3.65 | 0.17 | 0.247 | 1.90E-03 | 87.22 | 1.56 |

**Table 11.** Simulations with sparse genetic architecture and family structure. Mean and standard errors for effect estimates and test statistics of association tests at 1000 Null SNPs with true 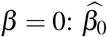 and 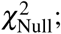; and at 1000 causal SNPs with an alternative hypothesis of true 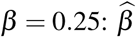 and *χ*^2^

| Model | mean($\beta_0$) | se($\beta_0$) | mean($\chi^2_{Null}$) | se($\chi^2_{Null}$) | mean($\beta$) | se($\beta$) | mean($\chi^2$) | se($\chi^2$) |
| --- | --- | --- | --- | --- | --- | --- | --- | --- |
| LR | 1.30E-03 | 1.60E-03 | 1.57 | 0.07 | 0.248 | 1.60E-03 | 47.46 | 0.73 |
| LR + 4 PCs | 1.20E-03 | 1.60E-03 | 1.53 | 0.07 | 0.248 | 1.60E-03 | 47.33 | 0.73 |
| LR + cvBLUP | 1.50E-03 | 9.00E-04 | 0.72 | 0.03 | 0.184 | 9.00E-04 | 35.81 | 0.49 |
| LMM | 2.00E-03 | 1.30E-03 | 0.98 | 0.05 | 0.249 | 1.30E-03 | 47.34 | 0.64 |
| BOLT-inf | 1.30E-03 | 9.00E-04 | 0.69 | 0.03 | 0.168 | 8.00E-04 | 33.39 | 0.44 |
| BOLT-LMM | 1.30E-03 | 9.00E-04 | 0.86 | 0.04 | 0.168 | 8.00E-04 | 64.13 | 0.79 |
| LR + true genetic effect | 1.70E-03 | 1.00E-03 | 1.14 | 0.05 | 0.250 | 1.00E-03 | 93.55 | 1.21 |

## References

Aulchenko, Y. S., De Koning, D.-J., and Haley, C. (2007). Grammar: a fast and simple method for genome-wide pedigree-based quantitative trait loci association analysis. Genetics.

Balding, D. J. and Nichols, R. A. (1995). A method for quantifying differentiation between populations at multi-allelic loci and its implications for investigating identity and paternity. Genetica, 96(1-2):3–12.

Benjamini, Y. and Yekutieli, D. (2001). The control of the false discovery rate in multiple testing under dependency. Annals of statistics, pages 1165–1188.

Burgess, S. and Thompson, S. G. (2013). Use of allele scores as instrumental variables for mendelian randomization. International journal of epidemiology, 42(4):1134–1144.

Chen, G.-B. (2014). Estimating heritability of complex traits from genome-wide association studies using ibs-based haseman–elston regression. Frontiers in genetics, 5:107.

Cortes, A., Dendrou, C. A., Motyer, A., Jostins, L., Vukcevic, D., Dilthey, A., Donnelly, P., Leslie, S., Fugger, L., and McVean, G. (2017). Bayesian analysis of genetic association across tree-structured routine healthcare data in the uk biobank. Nature genetics, 49(9):1311.

Dudbridge, F. (2016). Polygenic epidemiology. Genetic epidemiology, 40(4):268–272.

Gamazon, E. R., Wheeler, H. E., Shah, K., Mozaffari, S. V., Aquino-Michaels, K., Carroll, R. J., Eyler, A. E., Denny, J. C., Nicolae, D. L., Cox, N. J., et al. (2015). Predixcan: Trait mapping using human transcriptome regulation. bioRxiv, page 020164.

Gusev, A., Ko, A., Shi, H., Bhatia, G., Chung, W., Penninx, B. W., Jansen, R., De Geus, E. J., Boomsma, D. I., Wright, F. A., et al. (2016). Integrative approaches for large-scale transcriptome-wide association studies. Nature genetics, 48(3):245.

Hastie, T., Tibshirani, R., and Friedman, J. (2009). The elements of statistical learning: data mining, inference and prediction, page 244. Springer, 2 edition.

Kang, H. M., Sul, J. H., Service, S. K., Zaitlen, N. A., Kong, S.-y., Freimer, N. B., Sabatti, C., Eskin, E., et al. (2010). Variance component model to account for sample structure in genome-wide association studies. Nature genetics, 42(4):348.

Kang, H. M., Zaitlen, N. A., Wade, C. M., Kirby, A., Heckerman, D., Daly, M. J., and Eskin, E. (2008). Efficient control of population structure in model organism association mapping. Genetics, 178(3):1709–1723.

Khera, A. V., Chaffin, M., Aragam, K. G., Haas, M. E., Roselli, C., Choi, S. H., Natarajan, P., Lander, E. S., Lubitz, S. A., Ellinor, P. T., et al. (2018). Genome-wide polygenic scores for common diseases identify individuals with risk equivalent to monogenic mutations. Nature genetics, page 1.

Kolde, R. and Kolde, M. R. (2018). Package ‘pheatmap’.

Krapohl, E., Euesden, J., Zabaneh, D., Pingault, J., Rimfeld, K., Von Stumm, S., Dale, P., Breen, G., O’reilly, P., and Plomin, R. (2016). Phenome-wide analysis of genome-wide polygenic scores. Molecular psychiatry, 21(9):1188.

Laakso, M., Kuusisto, J., Stancakova, A., Kuulasmaa, T., Pajukanta, P., Lusis, A. J., Collins, F. S., Mohlke, K., and Boehnke, M. (2017). Metabolic syndrome in men (metsim) study: a resource for studies of metabolic and cardiovascular diseases. Journal of lipid research, pages jlr–O072629.

Lee, S. H., Yang, J., Goddard, M. E., Visscher, P. M., and Wray, N. R. (2012). Estimation of pleiotropy between complex diseases using single-nucleotide polymorphism-derived genomic relationships and restricted maximum likelihood. Bioinformatics, 28(19):2540–2542.

Lippert, C., Listgarten, J., Liu, Y., Kadie, C. M., Davidson, R. I., and Heckerman, D. (2011). Fast linear mixed models for genome-wide association studies. Nature methods, 8(10):833.

Listgarten, J., Kadie, C., Schadt, E. E., and Heckerman, D. (2010). Correction for hidden confounders in the genetic analysis of gene expression. Proceedings of the National Academy of Sciences, 107(38):16465–16470.

Liu, X., Mefford, J. A., Dahl, A., Subramaniam, M., Battle, A., Price, A. L., and Zaitlen, N. (2018). Gbat: a gene-based association method for robust trans-gene regulation detection. bioRxiv, page 395970.

Loh, P.-R., Tucker, G., Bulik-Sullivan, B. K., Vilhjalmsson, B. J., Finucane, H. K., Salem, R. M., Chasman, D. I., Ridker, P. M., Neale, B. M., Berger, B., et al. (2015). Efficient bayesian mixed-model analysis increases association power in large cohorts. Nature genetics, 47(3):284.

Maas, P., Barrdahl, M., Joshi, A. D., Auer, P. L., Gaudet, M. M., Milne, R. L., Schumacher, F. R., Anderson, W. F., Check, D., Chattopadhyay, S., et al. (2016). Breast cancer risk from modifiable and nonmodifiable risk factors among white women in the united states. JAMA oncology, 2(10):1295–1302.

Martin, A. R., Gignoux, C. R., Walters, R. K., Wojcik, G. L., Neale, B. M., Gravel, S., Daly, M. J., Bustamante, C. D., and Kenny, E. E. (2017). Human demographic history impacts genetic risk prediction across diverse populations. The American Journal of Human Genetics, 100(4):635–649.

McVean, G. (2009). A genealogical interpretation of principal components analysis. PLoS genetics, 5(10):e1000686.

Natarajan, P., Young, R., Stitziel, N. O., Padmanabhan, S., Baber, U., Mehran, R., Sartori, S., Fuster, V., Reilly, D. F., Butterworth, A., et al. (2017). Polygenic risk score identifies subgroup with higher burden of atherosclerosis and greater relative benefit from statin therapy in the primary prevention setting. Circulation, 135(22):2091–2101.

Nolte, I. M., van der Most, P. J., Alizadeh, B. Z., de Bakker, P. I., Boezen, H. M., Bruinenberg, M., Franke, L., van der Harst, P., Navis, G., Postma, D. S., et al. (2017). Missing heritability: is the gap closing? an analysis of 32 complex traits in the lifelines cohort study. European Journal of Human Genetics, 25(7):877.

Patterson, H. D. and Thompson, R. (1971). Recovery of inter-block information when block sizes are unequal. Biometrika, 58(3):545–554.

Patterson, N., Price, A. L., and Reich, D. (2006). Population structure and eigenanalysis. PLoS genetics, 2(12):e190.

Rakitsch, B., Lippert, C., Stegle, O., and Borgwardt, K. (2012). A lasso multi-marker mixed model for association mapping with population structure correction. Bioinformatics, 29(2):206–214.

Robinson, G. K. et al. (1991). That blup is a good thing: the estimation of random effects. Statistical science, 6(1):15–32.

Scutari, M., Mackay, I., and Balding, D. (2016). Using genetic distance to infer the accuracy of genomic prediction. PLoS genetics, 12(9):e1006288.

Seibert, T. M., Fan, C. C., Wang, Y., Zuber, V., Karunamuni, R., Parsons, J. K., Eeles, R. A., Easton, D. F., Kote-Jarai, Z., Al Olama, A. A., et al. (2018). Polygenic hazard score to guide screening for aggressive prostate cancer: development and validation in large scale cohorts. bmj, 360:j5757.

Svishcheva, G. R., Axenovich, T. I., Belonogova, N. M., van Duijn, C. M., and Aulchenko, Y. S. (2012). Rapid variance components–based method for whole-genome association analysis. Nature genetics, 44(10):1166.

Terry, R. B., Wood, P. D., Haskell, W. L., Stefanick, M. L., and Krauss, R. M. (1989). Regional adiposity patterns in relation to lipids, lipoprotein cholesterol, and lipoprotein subfraction mass in men. The Journal of Clinical Endocrinology & Metabolism, 68(1):191–199.

Torkamani, A., Wineinger, N. E., and Topol, E. J. (2018). The personal and clinical utility of polygenic risk scores. Nature Reviews Genetics, page 1.

Tucker, G., Loh, P.-R., MacLeod, I. M., Hayes, B. J., Goddard, M. E., Berger, B., and Price, A. L. (2015). Two-variance-component model improves genetic prediction in family datasets. The American Journal of Human Genetics, 97(5):677–690.

Vilhjálmsson, B. J., Yang, J., Finucane, H. K., Gusev, A., Lindström, S., Ripke, S., Genovese, G., Loh, P.-R., Bhatia, G., Do, R., et al. (2015). Modeling linkage disequilibrium increases accuracy of polygenic risk scores. The American Journal of Human Genetics, 97(4):576–592.

Warren, H., Casas, J.-P., Hingorani, A., Dudbridge, F., and Whittaker, J. (2014). Genetic prediction of quantitative lipid traits: comparing shrinkage models to gene scores. Genetic epidemiology, 38(1):72–83.

Wray, N. R., Goddard, M. E., and Visscher, P. M. (2007). Prediction of individual genetic risk to disease from genome-wide association studies. Genome research, 17(10):000–000.

Yang, J., Benyamin, B., McEvoy, B. P., Gordon, S., Henders, A. K., Nyholt, D. R., Madden, P. A., Heath, A. C., Martin, N. G., Montgomery, G. W., et al. (2010). Common snps explain a large proportion of the heritability for human height. Nature genetics, 42(7):565.

Yang, J., Lee, S. H., Goddard, M. E., and Visscher, P. M. (2011). Gcta: a tool for genome-wide complex trait analysis. The American Journal of Human Genetics, 88(1):76–82.

Yang, J., Zaitlen, N. A., Goddard, M. E., Visscher, P. M., and Price, A. L. (2014). Advantages and pitfalls in the application of mixed-model association methods. Nature genetics, 46(2):100.

Zaitlen, N., Kraft, P., Patterson, N., Pasaniuc, B., Bhatia, G., Pollack, S., and Price, A. L. (2013). Using extended genealogy to estimate components of heritability for 23 quantitative and dichotomous traits. PLoS genetics, 9(5):e1003520.

Zaitlen, N., Lindström, S., Pasaniuc, B., Cornelis, M., Genovese, G., Pollack, S., Barton, A., Bickeböller, H., Bowden, D. W., Eyre, S., et al. (2012a). Informed conditioning on clinical covariates increases power in case-control association studies. PLoS genetics, 8(11):e1003032.

Zaitlen, N., Paşaniuc, B., Patterson, N., Pollack, S., Voight, B., Groop, L., Altshuler, D., Henderson, B. E., Kolonel, L. N., Marchand, L. L., et al. (2012b). Analysis of case–control association studies with known risk variants. Bioinformatics, 28(13):1729–1737.

Zhou, X., Carbonetto, P., and Stephens, M. (2013). Polygenic modeling with bayesian sparse linear mixed models. PLoS genetics, 9(2):e1003264.

Zhou, X. and Stephens, M. (2012). Genome-wide efficient mixed-model analysis for association studies. Nature genetics, 44(7):821.

